# Protein complex prediction with AlphaFold-Multimer

**DOI:** 10.1101/2021.10.04.463034

**Authors:** Richard Evans, Michael O’Neill, Alexander Pritzel, Natasha Antropova, Andrew Senior, Tim Green, Augustin Žídek, Russ Bates, Sam Blackwell, Jason Yim, Olaf Ronneberger, Sebastian Bodenstein, Michal Zielinski, Alex Bridgland, Anna Potapenko, Andrew Cowie, Kathryn Tunyasuvunakool, Rishub Jain, Ellen Clancy, Pushmeet Kohli, John Jumper, Demis Hassabis

## Abstract

While the vast majority of well-structured single protein chains can now be predicted to high accuracy due to the recent AlphaFold [1] model, the prediction of multi-chain protein complexes remains a challenge in many cases. In this work, we demonstrate that an AlphaFold model trained specifically for multimeric inputs of known stoichiometry, which we call AlphaFold-Multimer, significantly increases accuracy of predicted multimeric interfaces over input-adapted single-chain AlphaFold while maintaining high intra-chain accuracy. On a benchmark dataset of 17 heterodimer proteins without templates (introduced in [2]) we achieve at least medium accuracy (DockQ [3] ≥ 0.49) on 13 targets and high accuracy (DockQ ≥ 0.8) on 7 targets, compared to 9 targets of at least medium accuracy and 4 of high accuracy for the previous state of the art system (an AlphaFold-based system from [2]). We also predict structures for a large dataset of 4,446 recent protein complexes, from which we score all non-redundant interfaces with low template identity. For heteromeric interfaces we successfully predict the interface (DockQ ≥ 0.23) in 70% of cases, and produce high accuracy predictions (DockQ ≥ 0.8) in 26% of cases, an improvement of +27 and +14 percentage points over the flexible linker modification of AlphaFold [4] respectively. For homomeric interfaces we successfully predict the interface in 72% of cases, and produce high accuracy predictions in 36% of cases, an improvement of +8 and +7 percentage points respectively.

## 1. Introduction

The formation of permanent and transient protein complexes underpins most biological processes, and understanding the structure of these complexes is a key step towards understanding and modifying their function. *In silico* protein structure prediction is playing an increasingly important role in biological research and applications as it has approached experimental accuracy, with the recent structure prediction system AlphaFold [1] providing a step change in structure prediction accuracy. Though AlphaFold was trained on individual protein chains, including many proteins whose structure was solved in complex with other proteins, it has shown remarkable ability to predict protein structures with bound co-factors or proteins stabilised by their multimeric interactions. Subsequent work has also shown that providing pseudo-multimer inputs (e.g. residue gap insertion or chains joined with a flexible linker) to the single-chain AlphaFold model is often successful at predicting multimer interactions [4, 5, 6, 7, 8, 9]. These papers have shown surprising generalization performance of the original trained AlphaFold model but leave open the question of how much more accurate AlphaFold is when training is also adapted for multimeric inputs.

The AlphaFold system has recently been described in publication [1] along with source code and model parameters [10]. To summarize briefly, it combines information from the amino acid sequence, multiple sequence alignments and homologous structures in order to predict the structure of individual protein chains. The core part of the neural network, called Evoformer, consists of a neural representation of the multiple sequence alignment (MSA) and pairwise relations between the different amino acids in the protein. These two representations are mixed and processed by a collection of neural network modules. The pair representation can be thought of as containing information about the relative positions of amino acids in the chain. This representation is used to predict the relative distances between the amino acids in the chain via a binned distance distribution (distogram). The first row of the MSA embedding is then used together with the pair embedding to predict the final structure. The model is trained end-to-end with gradients propagating from the predicted structure through the entire network.

In this work we extend AlphaFold to multiple chains during both training and inference, with native support for multi-chain featurization and symmetry handling. We refer to this system as AlphaFold-Multimer and demonstrate superior performance compared to existing approaches, including those based on using AlphaFold.

## 2. Methods

Multiple changes to the AlphaFold system were made to adapt it to training on protein complexes, which are detailed below. Summarizing briefly, we modify the losses to take into account permutation symmetry among identical chains, pair the MSA alignments between individual chains to surface cross-chain genetic information, introduce a new way of selecting subsets of residues for training, and make various small adjustments to the structure losses and the model architecture.

### 2.1. Multi-Chain Permutation Alignment

When computing the losses and scoring homomeric components, permutation symmetry must be taken into account. When a protein of a given sequence appears multiple times in the complex, the mapping between the predicted and the ground-truth coordinates is arbitrary and so the model cannot be assumed to predict chains in the same order as in the ground truth. To account for this we aim to pick the optimal permutation of predicted homomer chains that best matches the ground truth. The complexity of optimizing over all permutations grows combinatorially so we employ a simple heuristic that greedily tries to find a good permutation. The details of this heuristic are explained in subsection 7.3.

### 2.2. Multiple Sequence Alignment (MSA) Construction

Pairwise correlation between columns in an MSA has been shown to be a useful feature in predicting contacts and 3D structure. The per-chain MSAs used in AlphaFold-Multimer are constructed in a similar way to those used in AlphaFold. For homomeric complexes we simply copy the MSA *n* times, where *n* is the number of chain repeats, and stack these left-to-right. For heteromeric complexes where possible we pair sequences between the MSAs of each chain (see subsection 2.3) and stack these left-to-right. For partial alignments (where sequence pairings can be found between some chains but not others) we pad between the paired sequences with gap characters. The remaining unpaired MSA sequences for each chain are stacked in a block-diagonal fashion below the set of paired sequences, with padding on the off-diagonals. During training we bias sampling of the MSA cluster centres such that each chain’s unpaired MSA has an equal probability of being sampled from, regardless of the number of sequences it contains. Sampling of paired versus unpaired sequences is unbiased, such that the probability of selecting a paired versus unpaired row is proportional to their relative occurrence. At test time we achieve slightly better performance by performing unbiased MSA sampling, this may be because it yields the greatest diversity across recycling iterations.

### 2.3. Cross-Chain Genetics

Identifying pairwise correlation in an MSA requires prior determination and alignment of interacting homologs in the heteromeric case, which is in general ambiguous [11]. In this work we provide explicit aligned sequences to the network following the procedure of Zhou *et al*. [12] by pairing sequences using the UniProt species annotation. When multiple sequences exist for the same species we rank the candidate rows of each chain by similarity to their respective target sequence, and concatenate pairs of the same rank. More details are available in subsection 7.4.

### 2.4. Multi-Chain Cropping

The number of residues that AlphaFold-Multimer can be trained on is limited by memory and compute considerations as both memory use and compute use increase rapidly with the total number of amino acids in the complex. To ameliorate this, the AlphaFold system is trained on cropped segments of full length proteins, where these cropped regions are contiguous blocks of residues up to 384 residues.

We modified this procedure when training on complexes as the cropped regions need to involve multiple chains in a given complex, and binding interfaces between chains are crucial for modelling protein complexes. Therefore, we designed a cropping procedure that maximizes chain coverage and crop diversity whilst ensuring good balance between interface and non-interface regions. We balance per-chain contiguous cropping in sequence space with interface biased spatial cropping in coordinate space. The details of this procedure are described in subsection 7.2.

### 2.5. Architecture and Losses

AlphaFold uses a Frame Aligned Point Error (FAPE) loss [1], whereby the distances between ground truth and predicted atoms are computed in the local reference frame of every residue. In AlphaFold, this loss was clamped at 10 Å. For training the multimer model we made changes to the loss function used: for the intra-chain amino acid pairs of the complex we keep the same 10 Å clamping; for the inter-chain pairs we clamp the loss at 30 Å. This provides a better gradient signal for incorrect interfaces. Moreover, we add extra positional encodings denoting whether a given pair of amino acids corresponds to different chains and whether they belong to different homomer or heteromer chains.

To help prevent the model from predicting overlapping chains when it is highly uncertain we introduce an additional chain centre-of-mass loss term such that chains are pushed apart if they are closer than in the ground truth, defined as:

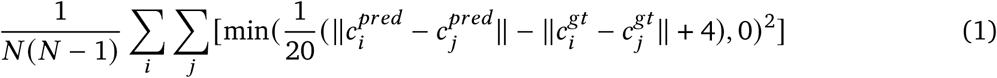

Where *N* is the number of chains, 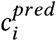 is the centre of mass of the C*α* atoms for predicted chain *i* and 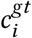 is the centre of mass of the C*α* atoms for ground truth chain *i*. The loss is clamped if the error is -4 Å or greater to prevent slight model uncertainty pushing the chains apart. The 1 20 weighting was chosen via a small hyper-parameter sweep.

In order to reduce the number of violations we made modifications to the violation losses from the original AlphaFold paper: we change the term penalizing steric clashes of non bonded atoms to average over clashing atoms rather than summing across all atoms. This is useful because initially all atoms will be at the origin, hence the gradient coming from clashes when summing will be extremely large early on in training, leading to instability. The clash loss in AlphaFold is modified as follows:

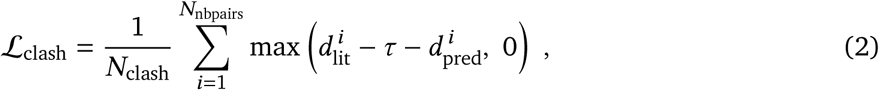

where 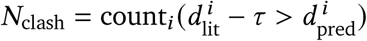 . Here, as in AlphaFold, the sum runs over nonbonded atom pairs, 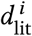 is the minimum distance between atoms derived from the relevant van der Waals radii. *τ* is a tolerance factor that is set to 1.5 Å. 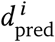 is the distance between the atoms in pair *i* in the predicted structure. To enable training with violation losses throughout training we lower the weight of the violation losses from 1. to 0.03 and lower the weight of the bond-angle term to ℒ_clash_ to 0.3. These factors were found by looking at the gradient magnitude of the different terms and setting values such that the gradients coming from violation losses would not dominate over the gradients coming from FAPE.

We also made various small changes to the model and implementation in order to facilitate inference of larger proteins for a given amount of memory. The details of these, and all other modifications mentioned in this section, are explained in the Supplementary Information.

### 2.6. Training Regimen

AlphaFold-Multimer was trained in a very similar way to AlphaFold [1]. The training dataset comprised structures from the Protein Data Bank (PDB) [13] with a maximum release date of 2018-04-30. Chains were sampled in inverse proportion to cluster size and their corresponding mmCIFs selected as input to the data pipeline, meaning that the rest of the chains in the bio-assembly will be included. This means that the probability of sampling a particular PDB entry is proportional to the sum of probabilities of the individual chain clusters for all chains in the file. The chain clusters are 40% identity clusterings of the Protein Data Bank with MMSeqs2 [14].

We randomly crop to 384 residues according to the sampling procedure outlined in subsection 2.4. We train the model to convergence (approximately 10M samples, for 2 weeks) across 128 TPUv3 cores with a batch size of 1 per TPU core. Then we fine-tune the model with half the learning rate, double the number of sequences fed into the MSA stack, and increase the violation loss weight (0.03->0.5) for two further days of training. We trained 2 models for the first stage; the best model on the validation set was then fine tuned with 5 different random seeds for the fine-tuning stage, producing 5 models in total.

### 2.7. Inference Regimen

At inference time we compare two different regimes. In the first we run all 5 trained models once and select the best model on each target according to model confidence, this is referred to as AlphaFold-Multimer (5×1). In the second regime we run each model 5 times with different random seeds, leading to 25 total predictions, this is referred to as AlphaFold-Multimer (5×5). Where it is not specified, AlphaFold-Multimer can be assumed to refer to AlphaFold-Multimer (5×5). The predictions are ranked using the model confidence described in subsection 2.8. The inference times for different sequence lengths are similar to those reported in [1].

### 2.8. Model Confidence

The AlphaFold model provides intrinsic model accuracy estimates in the form of a predicted TM-score (pTM) [1, §1.9.7]. We provide a similar metric for the AlphaFold-Multimer model, but modified to score interactions between residues of different chains, since we are predominantly interested in the accuracy of interfaces. We call this metric Interface pTM, or ipTM. See subsection 7.9 for more details.

In practice we use a weighted combination of pTM and ipTM as our model confidence metric, in order to account for some intra-chain confidence in our model ranking:

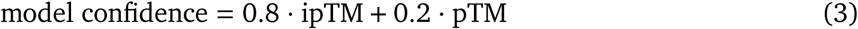

## 3. Related Work

Traditional approaches to multimeric structure prediction have tended to rely on a hybrid of template-based modelling and free docking [15, 16, 17, 18]. Template-based modelling approaches search for homologous complexes of known structure to serve as template models for the system under study. Free docking approaches start from individual subunits of known structure and attempt to select the most plausible interfaces from thousands of sampled orientations. The top 3 entrants of CASP14 Multimers — Baker-experimental [19], Venclovas [20] and Takeda-Shitaka lab [21] — all used methodologies based on these approaches. Takeda-Shitaka used a purely template-based approach, whilst Baker-experimental and Venclovas used a combination of template-based and free-docking. Baker-experimental additionally employed a deep learning based system to infer inter-chain contacts from co-evolution information, which were used to guide the overall sampling and scoring of docked structures.

Inspired by AlphaFold’s recent success at CASP14, there have now been a number of attempts to apply the trained AlphaFold network to the task of complex structure prediction. Many of these approaches simply add a gap or linker segment between chains of a complex, and then fold it with pre-trained AlphaFold as if it were a single chain [4, 5, 6, 7, 8, 9], the residue gap trick having been previously applied to RoseTTAFold [22]. The idea is that the neural network should identify the linker segment as unstructured and fold the single multi-domain chain in a similar way to multiple chains. Bryant *et al*. [9] benchmarked AlphaFold with a residue gap against RoseTTAFold and the rigid-docking method GRAMM [23], and AlphaFold substantially outperformed both other methods. Ghani *et al*. [2] combine this approach with docking using the ClusPro docking server [17]. Humphreys *et al*. [24] used AlphaFold with RoseTTAFold pre-screening to generate a dataset of 712 high confidence complexes, with previously unknown structure. All of these approaches use a single-chain trained AlphaFold or AlphaFold-like system at inference time; predictions on complexes are achieved simply by modifying the input. In this work we natively handle multimer chains at both training and inference time, and our system is trained on a dataset that includes multimers.

A key aspect of the AlphaFold model is its ability to learn to identify residue conservation and co-evolution directly from the MSA. Such an ability has also been shown to be a useful factor for predicting protein-protein interactions [25, 26]. However, constructing an informative multi-chain MSA is a challenging task, often involving the careful pairing of sequences between chains in order to construct valid interlogs of the target sequence that expose interface information. Various heuristic methods exist to perform this pairing [27, 28, 29, 12]. We follow the approach of Zhou *et al*. [12] by pairing by species label, disambiguating using similarity to the target sequence. Note that [5] only performs pairing for prokaryotes (using genetic distance), whilst [9] follows the same approach as ours and doesn’t distinguish between prokaryotes and eukaryotes, disambiguating in both cases with the similarity-based method.

## 4. Results

AlphaFold-Multimer predictions are generated as described in subsection 2.7. We benchmarked AlphaFold-Multimer on two datasets: *Benchmark 2* from [2], and *Recent-PDB-Multimers. Benchmark 2* is a set of 17 heterodimers from PDB after the training set cutoff date. The authors selected the targets such that there are no homologous complexes in PDB. *Recent-PDB-Multimers* is a homology-reduced set of 4,446 recent protein complexes from PDB. Predictions were made on full complexes before splitting into in-contact chain pairs (according to the ground truth) for analysis. These pairs were clustered to remove redundancy and filtered so that neither chain in the pair had greater than 40% template identity to the training set, yielding 2,609 unique pairs upon which we report metrics. For details see subsection 7.8. For both datasets we compare models using DockQ score [3, 30]. DockQ measures the quality of the interface and yields a score in the range [0, 1], interfaces with score *<* 0.23 are considered incorrect and scores *>* 0.8 are of the highest quality.

### 4.1. Benchmark 2

On *Benchmark 2* we compare AlphaFold-Multimer to the following systems:

- *AlphaFold-Linker*: In this setup we add a 21 residue repeated Glycine-Glycine-Serine linker between each chain before running it as a single chain through the AlphaFold model. Similar linker modifications to AlphaFold have previously been used [4].
- *AlphaFold-Gap (ColabFold [5])*: This is a third-party Google Colab, version from 2021-08-16, that runs AlphaFold with a 200 residue gap in the residue index between chains. It uses MMSeqs2 [14] for genetics, includes MSA pairing and doesn’t include templates.
- *ClusPro*: This setup runs AlphaFold on the individual chains before docking them together using ClusPro.
- *AlphaFold refined ClusPro*: The ClusPro predictions are refined by feeding them back into the AlphaFold model as templates. The resulting 10 predictions are re-ranked according to the interface predicted aligned error (PAE) score.
- *AlphaFold refined ClusPro plus AlphaFold*: This pools the *AlphaFold-Gap* predictions with those of *AlphaFold refined ClusPro* and re-ranks all 15 predictions using interface PAE.

For *ClusPro, AlphaFold refined ClusPro* and *AlphaFold refined ClusPro plus AlphaFold* we use the top 1 numbers from [2]. Figure 1a compares the average DockQ score between the different systems. The AlphaFold-Multimer system has an average DockQ Score of 0.63, while the ClusPro refined AlphaFold system has a score of 0.49. In Figure 1b we compare the relative difference in DockQ score between each system and AlphaFold-Multimer and estimate 95% confidence intervals using bootstrapping over the targets. A further breakdown of the results, and the per target DockQ scores are provided in Table S1 and Table S2.

**Figure 1.**
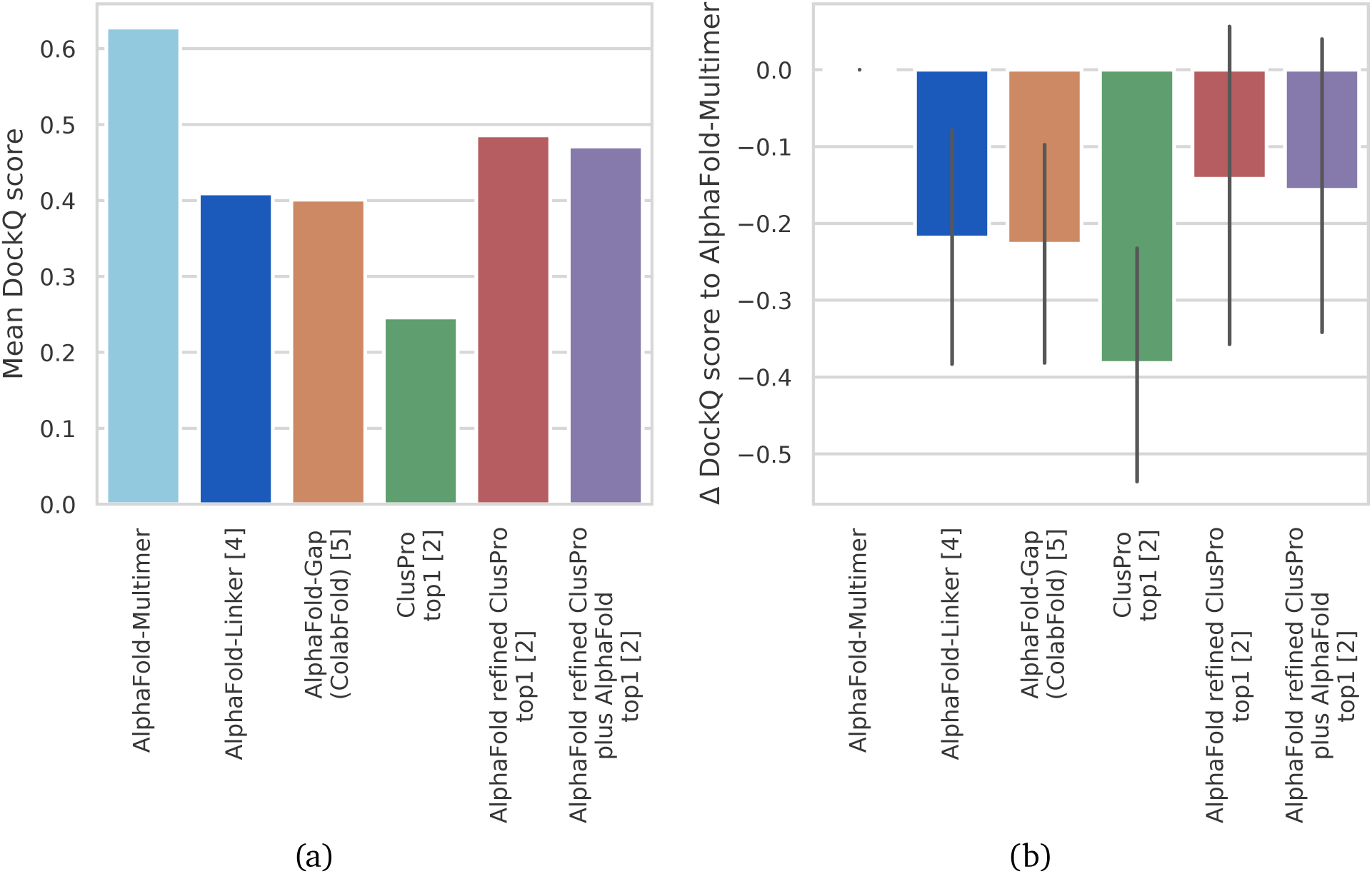
The performance of AlphaFold-Multimer against several published baselines is shown on the *Benchmark 2* dataset, consisting of 17 heterodimer targets with low training set homology. AlphaFold-Linker is AlphaFold with a 21 residue linker of repeated Glycine-Glycine-Serine residues, similar to previous AlphaFold modifications [4]. *AlphaFold-Gap (ColabFold [5])*, version from 2021-08-16, is a published system that runs AlphaFold with a gap between residue indices between chains, uses MMSeqs2 for genetics, includes MSA pairing and does not include templates. *ClusPro, AlphaFold refined ClusPro*, and *AlphaFold refined ClusPro plus AlphaFold* are all systems and results based on combining the docking algorithm ClusPro with AlphaFold, results are as reported in [2]. Error bars represent a 95% confidence interval around the mean.

### 4.2. Recent-PDB-Multimers

In Figure 2a we show the performance of the system for different levels of similarity to the training set on the *Recent-PDB-Multimers* dataset. Similarity is defined as identity × coverage for the least well-matched chain of the pair, maximised over all training set templates for which there is a match for both chains. Here we can see that there is fairly little correlation between similarity to the training set and model quality.

**Figure 2.**
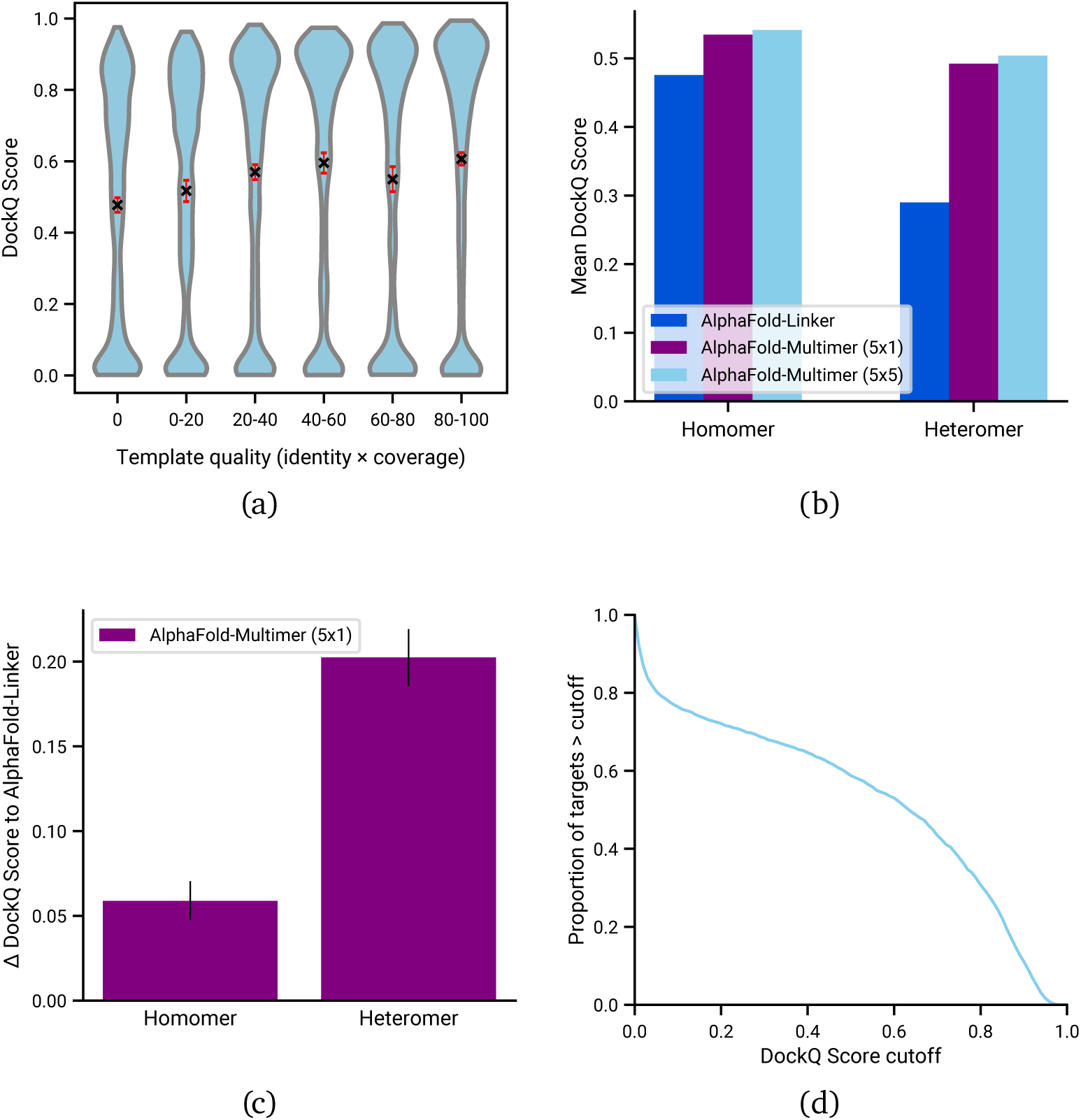
The performance of AlphaFold-Multimer on the *Recent-PDB-Multimers* dataset: the structures of 4,446 non-redundant protein complexes from recent PDB were predicted, from which all non-redundant pairs in contact were extracted (defined as any heavy atom of one chain being within 5A of any heavy atom of the other chain), yielding 5,287 pairs. For figures b, c and d this set was further reduced such that no chain in the pair had greater than 40% template overlap (identity × coverage) to the training set, yielding 1,286 homomeric and 1,323 heteromeric scored interfaces. All error bars represent a 95% confidence interval around the mean.

In Figure 2b we compare the average DockQ score between AlphaFold-Multimer (5×1) and AlphaFold-Linker. We see that AlphaFold-Multimer (5×1) is more accurate on both homomeric and the heteromeric interfaces: in the homomeric case the improvement is relatively modest with +0.06 mean DockQ score, but in the heteromeric case the improvement is more pronounced with +0.2 mean DockQ score. On average the performance on homomeric interfaces is better than for heteromeric ones. We note that increasing the number of inferences per model to 5 (AlphaFold-Multimer (5×5)) yields a further small gain of around +0.01 DockQ. In Figure 2c we show the difference between the two systems with bootstrapped error bars, showing clearly that AlphaFold-Multimer demonstrates a statistically significant improvement. The proportion of targets above different DockQ scores is shown in Figure 2d, a further breakdown into DockQ categories (Incorrect, Acceptable, Medium and High) is available in Table S3. The AlphaFold-Multimer system successfully predicts (DockQ ≥ 0.23) heteromeric interfaces in 70% of cases, and successfully predicts homomeric interfaces in 72% of cases.

In Figure 3 we show the distribution of DockQ score as a function of predicted interface TM-score for the AlphaFold-Multimer system. The figure shows that our predicted accuracy metric is a good predictor of model quality with the majority of high confidence predictions also having high accuracy.

**Figure 3.**
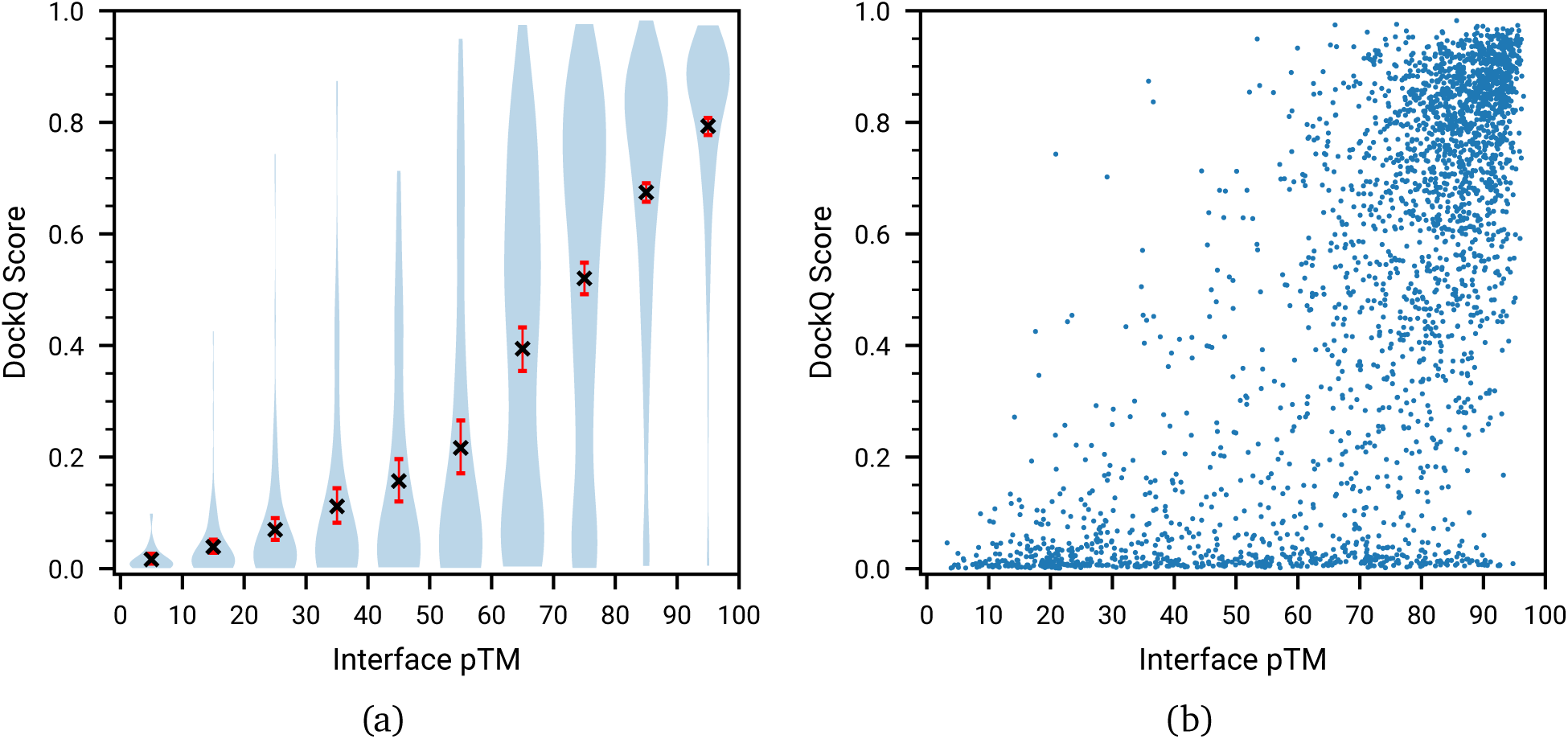
The correlation between AlphaFold-Multimer model confidence (predicted interface TM-score) and the true DockQ score is shown for the 2,648 homology reduced in-contact chain pairs from the *Recent-PDB-Multimers* dataset. Error bars represent a 95% confidence interval around the mean. In (a) interface pTM is grouped into intervals of size 10, (b) shows the raw data points.

In Figure 4 we show a number of examples where AlphaFold successfully predicted the right multimeric structure. In order to illustrate how to understand the predicted error metric we show an example of a case with incorrect coordinates (PDB code: 6QF7) which is also predicted to be wrong in Figure 5. In this case the individual chains are predicted to be correct, while their predicted relative position is uncertain, as can be seen from the blocks with a high predicted error.

**Figure 4.**
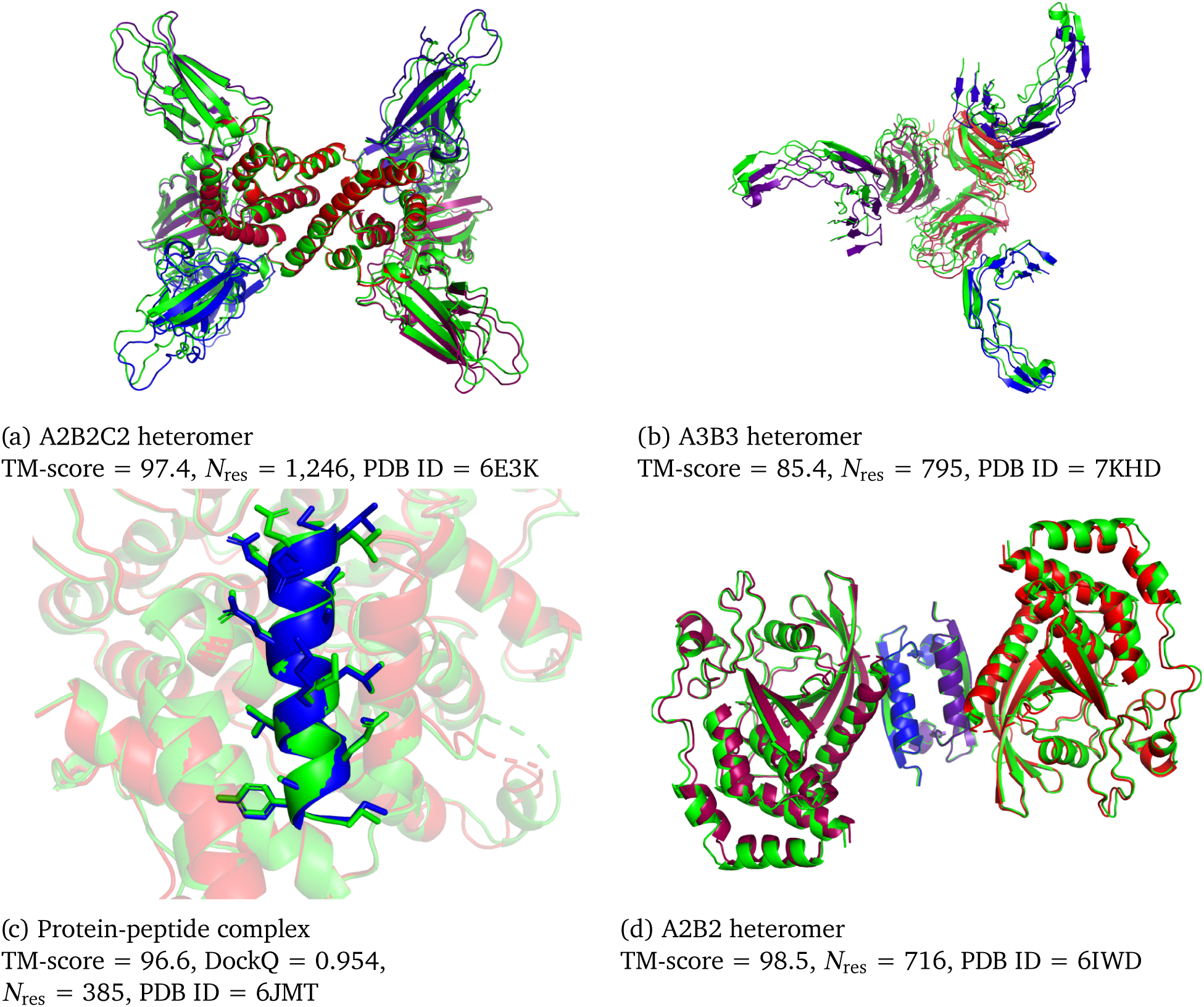
Structure examples predicted with the AlphaFold-Multimer. Visualised are the ground truth structures (green) and predicted structures (coloured by chain).

**Figure 5.**
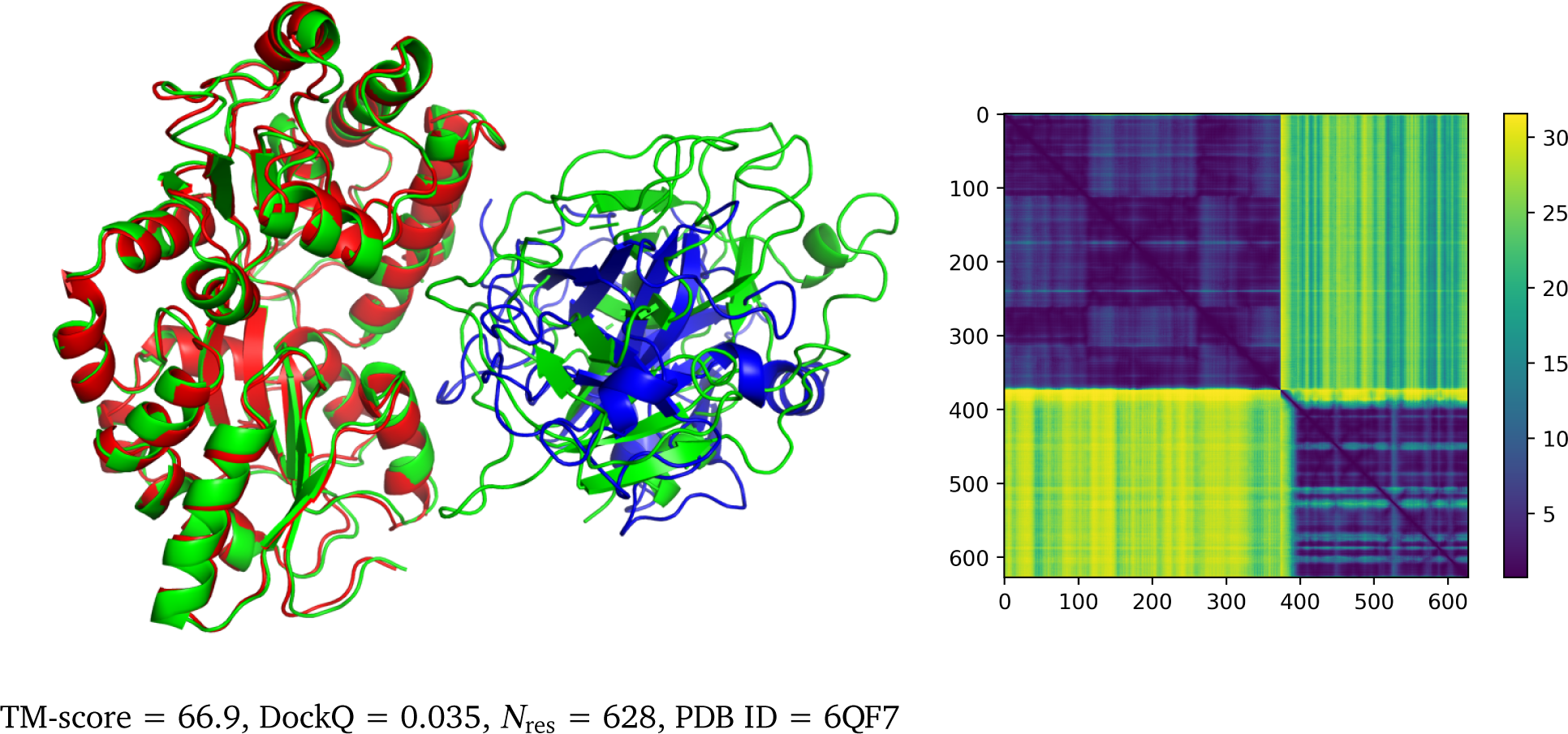
Example of a predicted heterodimer with incorrect geometry that is correctly predicted as low confidence by the predicted aligned error (PAE). Visualised are the ground truth structures (green), predicted structures (coloured by chain), and PAE heat map. The PAE heat map shows the predicted error (in Angstroms) between all pairs of residues.

The AlphaFold system was trained to predict the individual chains of multimeric complexes, and as such the model was not given the context provided by the other chains explicitly. Nevertheless AlphaFold has been shown to often implicitly model structures with missing context correctly. In the AlphaFold paper [1] an example is provided of a heavily intertwined multimer with correctly predicted but not physically meaningful chain structure. This leads us to ask the question whether the individual chain predictions of AlphaFold-Multimer are more accurate than the corresponding AlphaFold predictions. In Figure 6 we compare the performance of AlphaFold-Linker to AlphaFold-Multimer under two regimes. In AlphaFold-Multimer (5×1) the full complex is folded before extracting the individual chains, so that the system sees additional context. In the second regime (*AlphaFold-Multimer as Monomers* and *AlphaFold*) the individual chains are folded in isolation. We can see that in the case of heteromers AlphaFold-Multimer is less accurate than AlphaFold when given the single chains with incorrect monomer stoichiometry, but is more accurate than AlphaFold when the single chains are predicted as part of the full complex. We do not observe a similar effect in the AlphaFold-Linker system.

**Figure 6.**
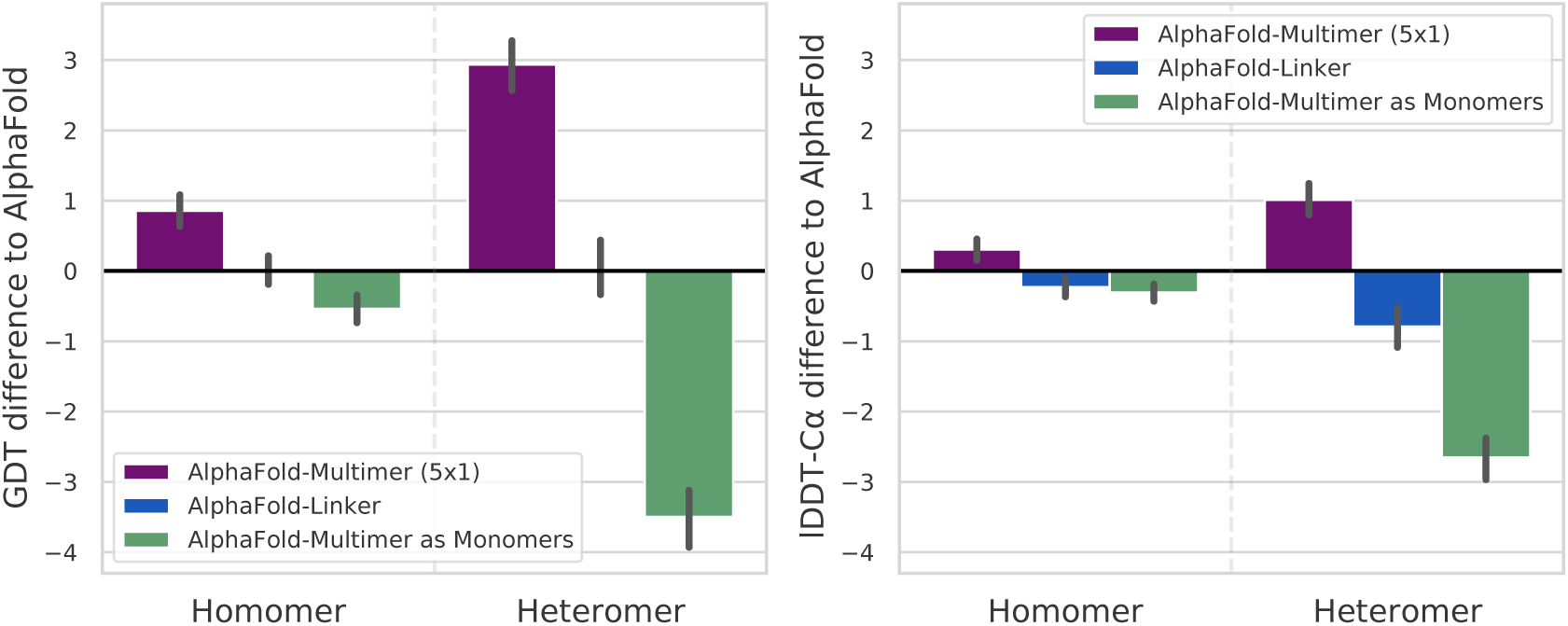
Comparison on 5,091 clustered single chains extracted from the *Recent-PDB-Multimers* dataset. Metrics for AlphaFold-Linker and AlphaFold-Multimer (5×1) are computed by predicting the full-complex, then cropping out the individual chains. For *AlphaFold* and *AlphaFold-Multimer as Monomers* the individual chains are run with A1 stoichiometry instead of the true stoichiometry. All error bars represent a 95% confidence interval around the mean.

## 5. Discussion

We have shown that by modifying the AlphaFold architecture to natively handle multimers and training explicitly on oligomeric data, we are able to provide high accuracy predictions for a large fraction of PDB complexes, surpassing the accuracy of inference-only modifications to AlphaFold. We have not yet implemented multimer templates or self-distillation of multimer predictions, so there is also likely substantial scope for future accuracy improvements.

We observe that performance is generally higher for homomeric interfaces than for heteromeric interfaces; this is presumably since in the homomeric case the MSA will readily encode evolutionary information about the complex interfaces, while this information is more limited and harder to access in the case of heteromeric interfaces. As a limitation, we observe anecdotally that AlphaFold-Multimer is generally not able to predict binding of antibodies and this remains an area for future work. We have also shown that the confidence metrics provided by the model correlate well with the true accuracy, something that is vital for the useability of a structure prediction model. By allowing accurate prediction of protein complexes we hope that this method will enable biologists to further accelerate the recent progress in structural bioinformatics.

The source code, weights and inference scripts for the trained AlphaFold-Multimer model are available at https://github.com/deepmind/alphafold.

## 6. Acknowledgements

We thank C. Beattie, H. Maclean, A. Huber, C. Meyer, C. Donner, A. Ballard, M. Figurnov, S. Nikolov, Z. Wu, J. Adler, J. Dunger, O. Vinyals, F. Yang and all our colleagues at DeepMind, Google and Alphabet for their encouragement and support.

## 7. Supplementary Information

### 7.1. Data

This section describes the AlphaFold-Multimer data pipeline, which generates features for the model. The input to the data pipeline is an mmCIF file containing one or more chains. The AlphaFold-Multimer data pipeline follows similar steps to the single-chain AlphaFold system [1], with a few key differences that are described below.

We derive the details about the structure’s biological assemblies from the mmCIF file and uniformly sample a single bio-assembly from all the available bio-assemblies. The MSA features are merged using the method outlined in subsection 2.2.

The AlphaFold-Multimer system uses only per-chain templates. The template search is similar to the AlphaFold system template search, except that it uses HMMER suite v3.3 hmmsearch and hmmbuild instead of HHsearch. First, the Uniref90 MSA obtained in the MSA search is converted to an HMM using hmmbuild (the only flag set to a non-default value is –hand). HMMER hmmsearch is then used to search for matches of the HMM against pdb_seqres.txt, downloaded from ftp://ftp.wwpdb.org/pub/pdb/derived_data/pdb_seqres.txt on 2020-05-14. We limit the number of templates to 20. Further processing is as described in the AlphaFold paper. Any structure released after 2018-04-30 is excluded from training. The HMMSearch flags used are `–F1 0.1 –F2 0.1 –F3 0.1 –incE 100 -E 100 –domE 100 –incdomE 100`.

The training data rebalancing procedure is very similar to the AlphaFold single chain setup: we uniformly sample a chain cluster, then uniformly sample a chain within that cluster and select the mmCIF to which it belongs as input to the pipeline. The chain clusters are 40% identity clusterings of the Protein Data Bank with MMSeqs2 [14].

During the training procedure we use a different self-distillation set, set of filters, MSA preprocessing, and residue cropping methods from the AlphaFold data pipeline. The self-distillation set is created using the clustered MGnify dataset [31], filtered to clusters with more than 10 sequences, resulting in about 13M sequences.

We filter out proteins during training if:

- The input mmCIF has resolution greater than 9 Å.
- Any single amino acid accounts for more than 80% of the complex sequences.
- The example comes from the distillation set and has less than 200 residues.

Features of the individual chains are merged before further processing. For all identical chains, the features are concatenated along the residue dimension. *N*_res_ × *N*_seq_ features of different chains are paired using the cross-chain genetics method outlined in subsection 7.4. *N*_res_ features were simply concatenated along the residue dimension.

The merged features are processed in a similar manner to AlphaFold, including MSA sampling and clustering, apart from the differences outlined in subsection 2.2. MSA block deletion is not included in the AlphaFold-Multimer data pipeline.

The features input to the multimer model are identical to AlphaFold with three extra features:

- asym_id - a unique integer per chain indicating the chain number. The ordering of the input chains is arbitrary as the network is agnostic to chain order.
- entity_id - a unique integer for each set of identical chains, for example in an A3B2 stoichiometry complex all the “A” chains would have the same ID and all the “B” chains would have the same ID.
- sym_id - a unique integer within a set of identical chains, for example in an A3B2 stoichiometry complex the “A” chains would have IDs [0, 1, 2] and the “B” chains would have IDs [0, 1].

### 7.2. Multi-chain Cropping

#### 7.2.1. Contiguous Cropping

In this method (Algorithm 1), we iterate through the list of chains, selecting a contiguous crop from each until we have reached our *N*_res_ budget. We keep track of two variables during the procedure: *n*_added_ is the number of residues selected so far, and *n*_remaining_ is the combined length of yet to be cropped chains (excluding the current chain). crop_size_min taken for each chain ensures we use as much of our *N*_res_ budget as possible, and crop_size_max ensures we don’t go over it. The input to the algorithm is the set of chain lengths, {*n*_*k*_} , and the total residue budget *N*_res_. The output is a set of masks to apply to each chain in order to extract the corresponding crops, {m_*k*_} . The chains are first randomly shuffled to avoid bias.

##### Algorithm 1 Crop Contiguous

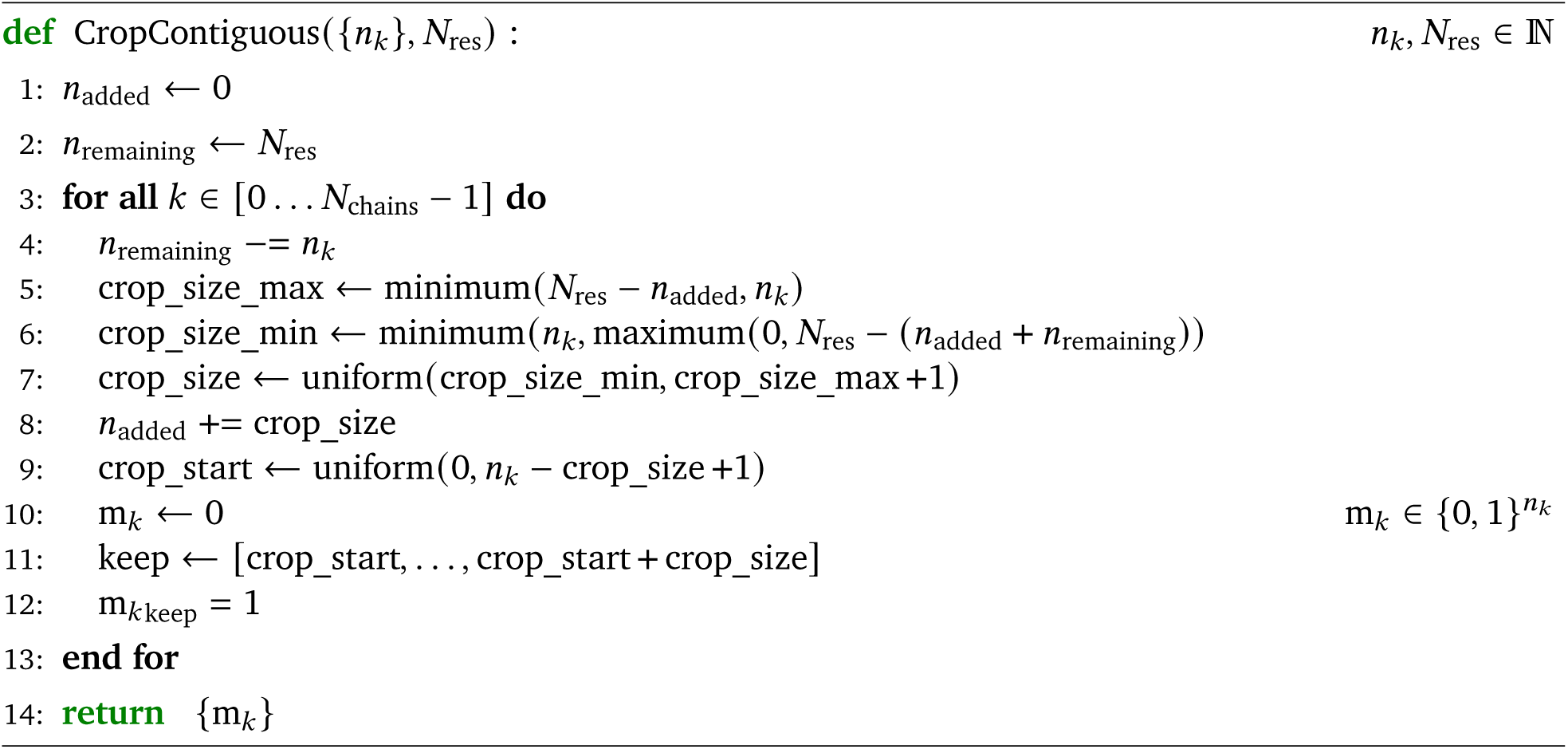

#### 7.2.2. Spatial Cropping (Interface Biased)

In this method (Algorithm 2), we specifically target interface regions by selecting a 3D spatial crop around a randomly selected interface residue. The crop consists of the *N*_res_ spatially nearest neighbours to the selected interface residue, as defined by distances between C*α* coordinates. The input to the algorithm is the set of C*α* coordinates across all chains, {x_*i*_}, and the index of the selected interface residue, *c*. The output is a set of binary values indicating which residues to include in the crop, {*m*_*i*_}. Distance ties are broken by adding small uniquifying values. We found empirically that a 50:50 ratio of interface cropping to contiguous cropping worked best for multimers (always contiguous cropping for monomers).

### 7.3. Multi-Chain Permutation Alignment

Stoichiometry must be carefully accounted for when scoring multimer structure predictions. In a prediction for A2B, both orderings of the A chains are equally valid, regardless of their ordering in the ground truth. In particular, if this is not accounted for in the loss then correct predictions will be unfairly penalized and the network will fail to train properly.

#### Algorithm 2 Crop Spatial

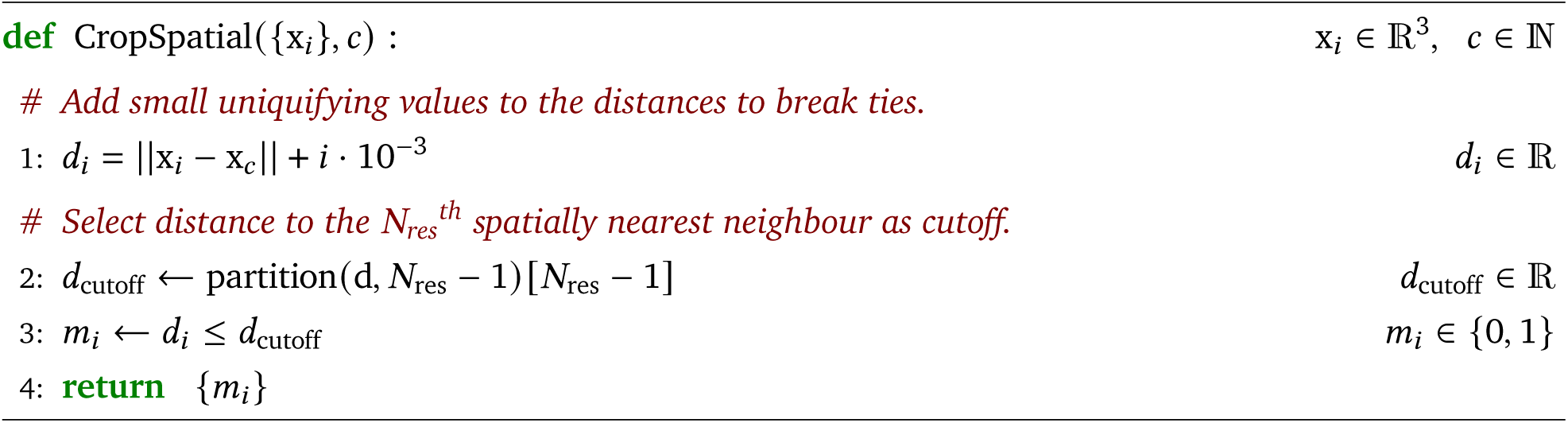

To address this issue, before scoring a multimer prediction we first permute chains with identical sequences such that they are best-effort aligned with those of the prediction. One could imagine considering the best alignment over all possible permutations, however this quickly becomes intractable, so a heuristic method is needed. Our method is a simple greedy heuristic that can be performed efficiently on TPU; other multi-chain alignment algorithms also exist [32, 33]. The optimal ground truth chain permutation is found using Algorithm 3, which can be broken down into the following alignment and assignment stages.

#### 7.3.1. Alignment Stage

In the alignment phase, we pick a pair of anchor chains to align, one in the ground truth and one in the prediction. The ground truth anchor chain *a*^gt^ is chosen to be the least ambiguous possible, for example in an A3B2 complex an arbitrary B chain is chosen. In the event of a tie e.g. A2B2 stoichiometry, the longest chain is chosen, with the hope that in general the longer chains are likely to have higher confident predictions. The prediction anchor chain is chosen from the set 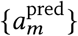 of all prediction chains with the same sequence as the ground truth anchor chain. We then transform the ground truth chain C*α* coordinates 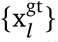 onto the prediction chain C*α* coordinates 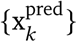 such that the chosen pair of anchor chains is optimally aligned (line 2).

#### 7.3.2. Assignment Stage

In the assignment stage (Algorithm 4), we greedily assign each of the predicted chains to their nearest neighbour of the same sequence in the ground truth. These assignments define the optimal permutation to apply to the ground truth chains. Nearest neighbours are defined as the chains with the smallest distance between the average of their C*α* coordinates.

We repeat the above alignment and assignment stages for all valid choices of the prediction anchor chain 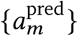 given the ground truth anchor chain *a*^gt^. Finally, we pick the permutation (line 8) that minimizes the RMSD between the C*α* coordinate averages of the predicted and ground truth chains.

The above procedure must be modified during training to account for cropping. In particular, in the alignment stage, we must first restrict the ground truth anchor chain to the same crop region as the paired prediction chain before alignment. Similarly, in the assignment stage, for each predicted chain the ground truth chains must be restricted to the same crop region before assignment.

##### Algorithm 3 Multi-chain permutation alignment

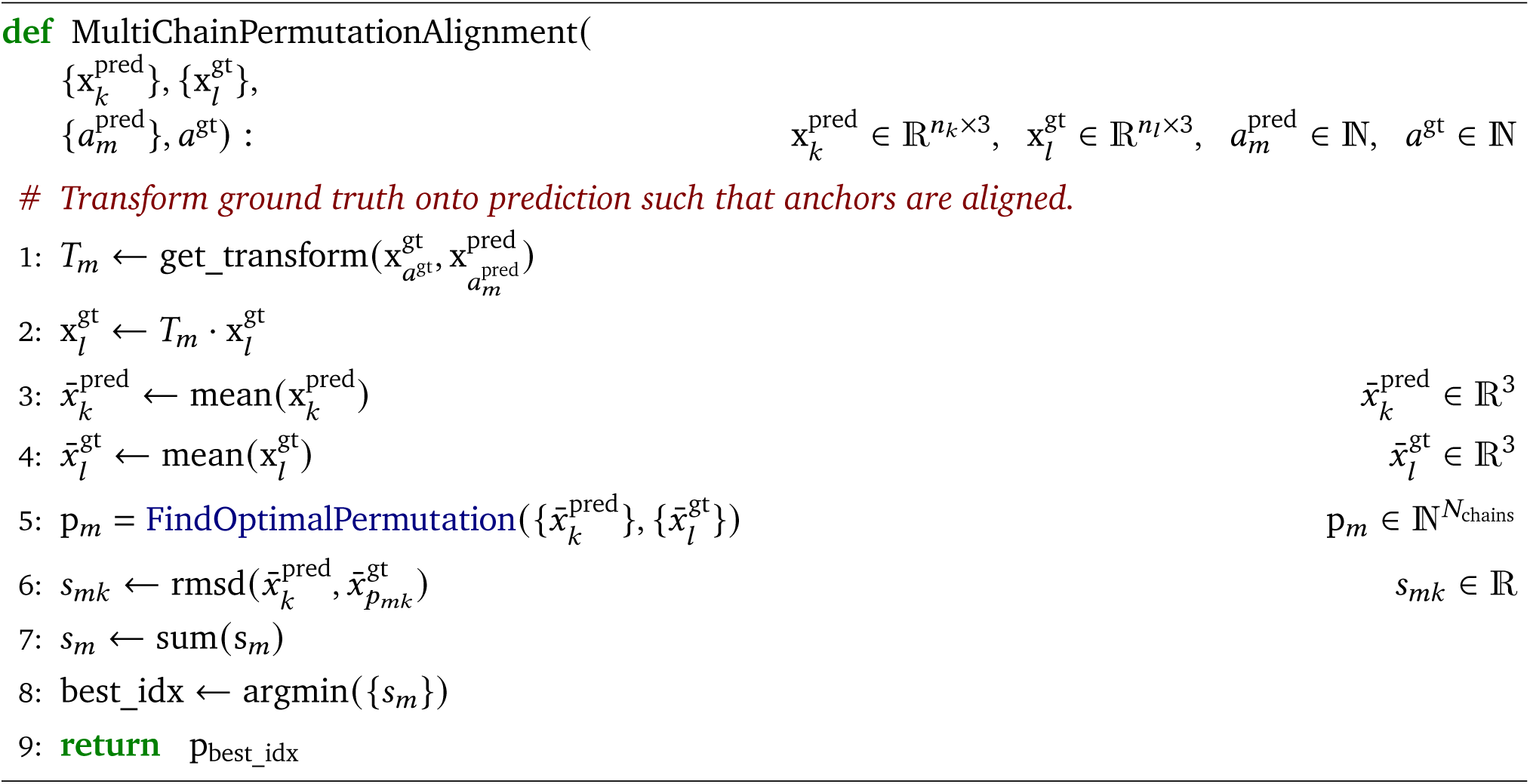

##### Algorithm 4 Find Optimal Permutation

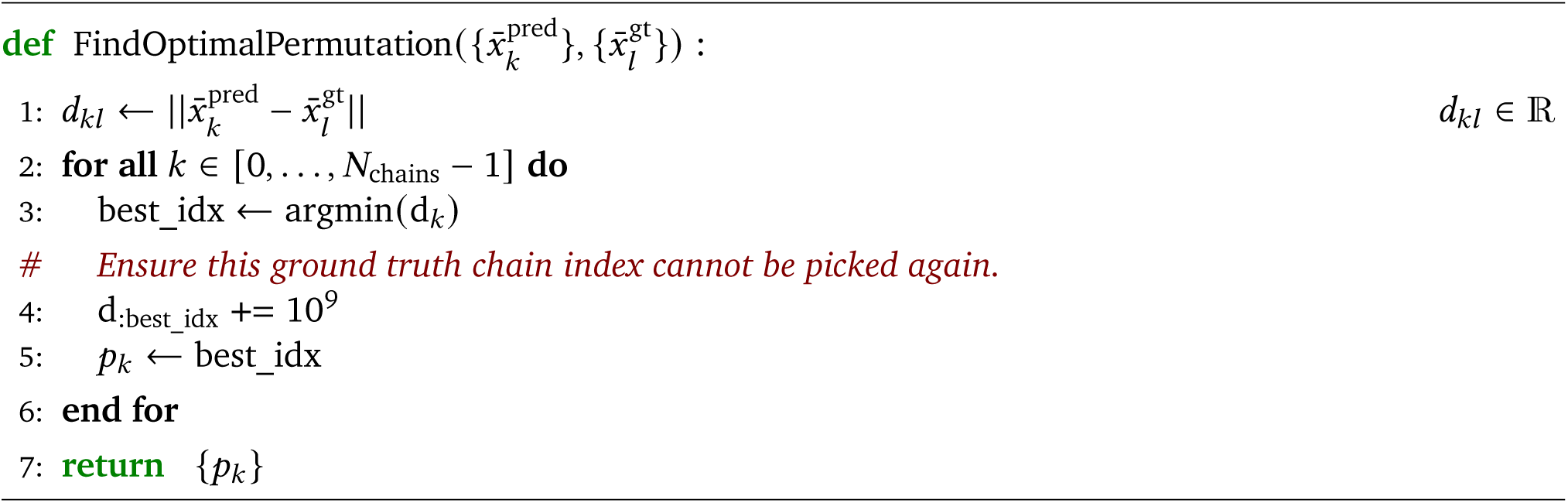

### 7.4. Cross-Chain Genetics

In order to make cross-chain coevolutionary information available to the network, it is necessary to pair sequences between the per-chain MSAs such that they form valid orthologs of the target sequence. In practice, this pairing can’t be done exactly, and various heuristic methods are used instead to approximate it. We follow the procedure of Zhou *et al*. [12] and pair sequences of the same species and disambiguate according to sequence similarity:

1. For each chain, JackHMMER v3.3 [34] is used to query the unclustered UniProt 2020_05 database [35] with maximum MSA depth set to 50,000, and non-default flags values set to -N 1 -e 0.0001 -Z 119222328.
2. The per-chain MSA sequences are grouped by species, using species labels derived from UniProt’s idmapping downloaded from https://ftp.uniprot.org/pub/databases/uniprot/current_release/knowledgebase/idmapping/ for the 2020_05 UniProt version.
3. Sequences are then paired within a specific species. If there is only one sequence per species, we simply concatenate them. Otherwise, we need to disambiguate.
4. We match the chain MSAs by ordering them by sequence identity to the target sequence and matching sequences of the same rank. There will be as many matches as there are hits in the shortest MSA.
5. Chains with empty MSAs are not matched and the resulting paired MSA will be padded with gap (no-match) symbols.

### 7.5. Chain Relative Positional Encoding

In AlphaFold [1], relative positional features are encoded into the initial pair representation of the network, allowing the network to reason about the positions of residues in the chain and break symmetry between identical residues. In the case of multimers, we also need to tell the network when two residues are in different chains, allowing it to break symmetry between homomer chains. To do this (Algorithm 5) we simply add an extra bin to the relative position encoding 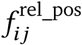 of AlphaFold to denote ‘different chain’. Furthermore, we also provide a feature to the network 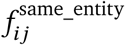 encoding whether the residues are from chains with identical sequences, and another feature 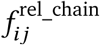 encoding relative chain index between chains of identical sequence. Relative residue indices are clipped between [−*r*_max_, *r*_max_], with *r*_max_ = 32, and relative chain indices are clipped between [−*s*_max_, *s*_max_], with 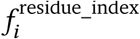 is the index of residue *i*, and 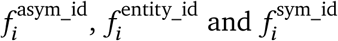 are chain identifiers for residue 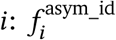 distiguishes between chains, 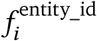 distinguishes between unique chain sequences, and 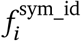 distinguishes between chains of the same sequence.

#### Algorithm 5 Relative position encoding

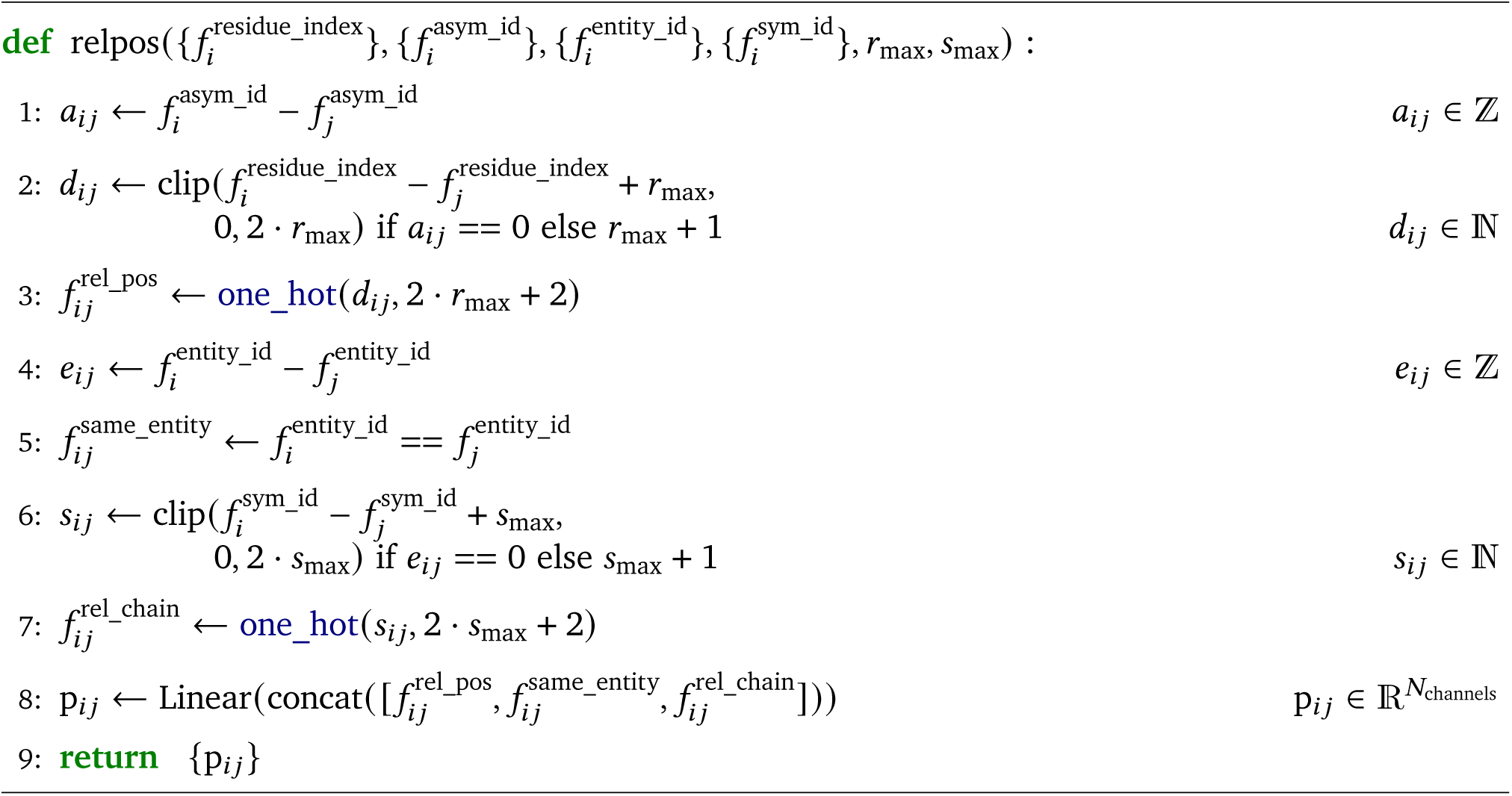

### 7.6. Mixed Frame Aligned Point Error (FAPE) Clamping

In AlphaFold the frame aligned point error (FAPE) backbone loss is clamped at 10 Å, emphasizing accurate positioning of residues within their local neighbourhood, but de-emphasizing longer range errors. For multimer chains, we also want to encourage accurate positioning of interface residues, which are in general much farther apart than neighbouring within-chain residues. We achieve this by splitting the FAPE loss into two. The first is clamped at 10 Å and applied to within-chain residue-pairs; the second is clamped at 30 Å and applied to between-chain residue-pairs. These two losses are then summed (which also has the effect of upweighting between-chain residue-pairs).

#### Algorithm 6d One-hot encoding with nearest bin

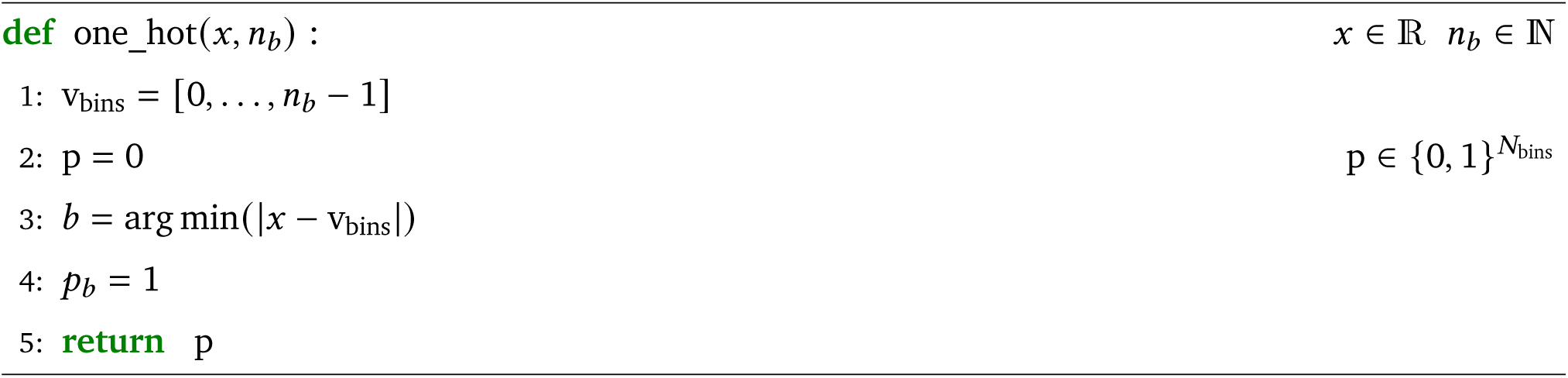

### 7.7. Architectural Modifications

We made various small architecture changes to facilitate inferencing proteins of larger size, and made some further minor changes to make different components of the model more consistent with one another.

We swap the order of the attention and triangular multiplicative update layers in the template stack relative to the AlphaFold model. This is to make it consistent with the order in the Evoformer stack. The template unit vectors that were disabled in the AlphaFold model are enabled here, i.e. they are not set to zero but computed from the template coordinates. We changed the way the template embeddings are aggregated. Before the model would give a pairwise representation of embeddings for each template, which were then aggregated via attention independently for each pair of residues. Here we use a simpler approach whereby we simply average the template embeddings. For efficiency we compute this similarly to how a moving average would be computed: we sum the embeddings as they are getting produced and divide by number of templates at the end. This has the advantage that it does not require template embeddings for all templates in order to average them, which leads to a substantial reduction in memory consumption. We have observed no significant drop in accuracy when aggregating templates in this way. Furthermore we moved the outer product mean to the start of the Evoformer block. This allows the pair representation and the MSA representation to evolve independently within a given block, with all cross communication happening at the start of the block. At training time this means that we do not need to keep all of the MSA stack activations and all the pair stack activations in memory at the same time during backpropagation. At inference time both stacks can be processed in parallel.

### 7.8. Datasets

The *Recent-PDB-Multimers* set consists of all targets in the Protein Data Bank [13] released between 2018-04-30 and 2021-08-02. This set was filtered to proteins with more than one chain, less than 9 chains and less than 1,536 total residues. It was also clustered with the following approach, which yields a set of 4,446 protein complexes:

1. Assign each chain to its 40% overlap cluster (using the clusters provided by PDB (link)).
2. Assign each protein complex a cluster identifier that is the union of all the single-chain cluster ids from step 1.
3. Randomly pick a single protein complex from every full-complex cluster.

Predictions were made on the full complexes, however for *Recent-PDB-Multimers* metrics were computed for each chain-pair separately. The chain pairs were selected if they were ‘in contact’ in the ground truth, defined as any heavy atom of one chain being within 5A of any heavy atom of the other chain. They were then clustered to remove redundancy: chain pairs were greedily selected if the cluster id pair was unique amongst those already selected. Finally the data was filtered such that no chain has greater than 40% template identity to the training set. This resulted in 2,609 unique in-contact chain-pairs (5,287 pre-filtering).

### 7.9. Interface pTM

Defining an interface version of AlphaFold’s predicted TM-score metric [1, §1.9.7] amounts to modifying [1, Equation 40] so that *i* and *j* come from different chains.

Concretely, if we follow the notation in [1, §1.9.7] such that 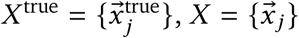 are the C*α* positions of the ground truth and predicted structures respectively, 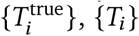 are the corresponding backbone frames, 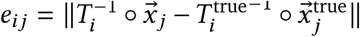 is the error in the position of the C*α* atom of residue *j* when the predicted and true structures are aligned using the backbone frame of residue *i, N*_res_ is the total number of residues, and 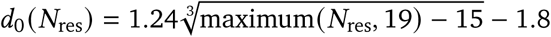 is the TM-score normalization constant, then we can obtain a interface TM-score prediction as:

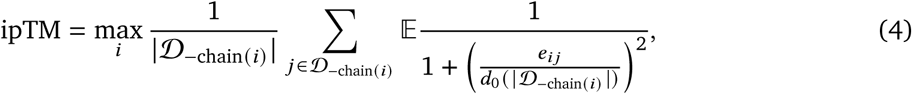

where 𝒟_chain(*i*)_ is the set of all residues except those of the chain of residue *i*. As in [1, §1.9.7], the expectation is taken over the probability distribution defined by *e*_*i,j*_ , which is predicted by a neural network.

**Table S1.**
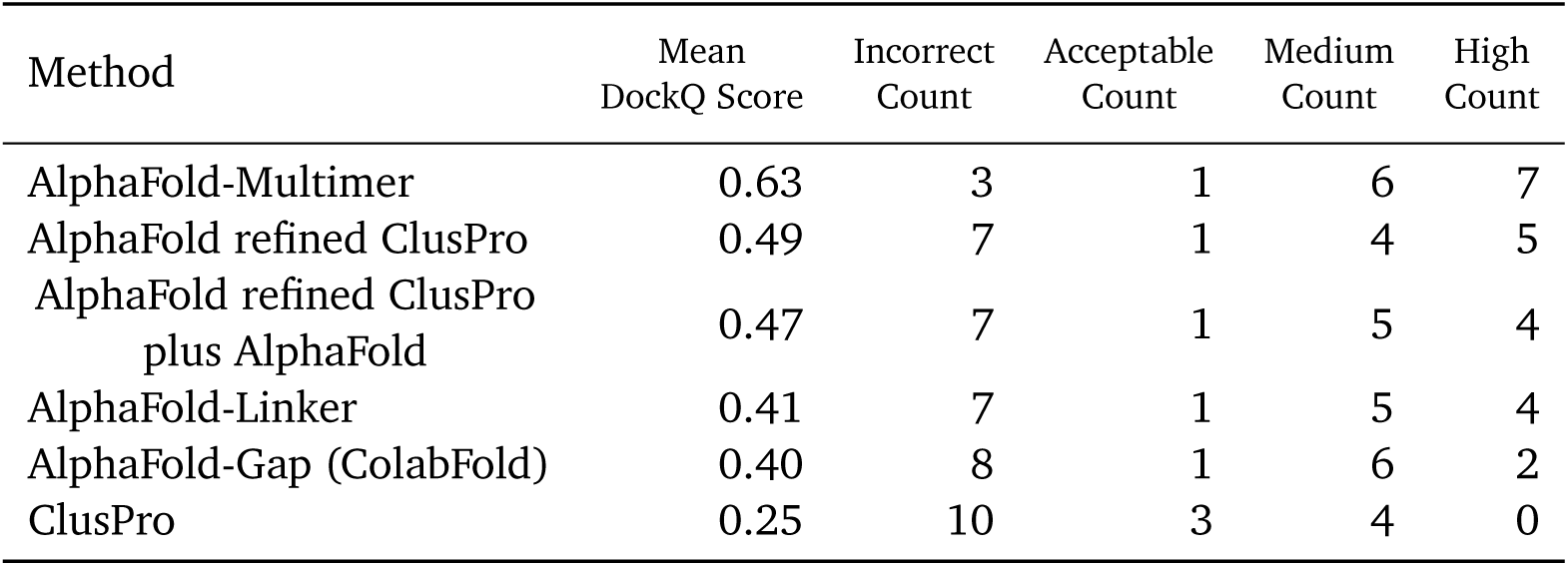
Performance on the Benchmark 2 dataset from [2] consisting of 17 heterodimers with low training set similarity. The following CAPRI definitions were used: Incorrect: 0 ≤ DockQ *<* 0.23 Acceptable: 0.23 ≤ DockQ *<* 0.49 Medium: 0.49 ≤ DockQ *<* 0.80 High: 0.80 ≤ DockQ Target DockQ Score 5ZNG 0.69

| Method | Mean DockQ Score | Incorrect Count | Acceptable Count | Medium Count | High Count |
| --- | --- | --- | --- | --- | --- |
| AlphaFold-Multimer | 0.63 | 3 | 1 | 6 | 7 |
| AlphaFold refined ClusPro | 0.49 | 7 | 1 | 4 | 5 |
| AlphaFold refined ClusPro plus AlphaFold | 0.47 | 7 | 1 | 5 | 4 |
| AlphaFold-Linker | 0.41 | 7 | 1 | 5 | 4 |
| AlphaFold-Gap (ColabFold) | 0.40 | 8 | 1 | 6 | 2 |
| ClusPro | 0.25 | 10 | 3 | 4 | 0 |

**Table S2.** Results per target for AlphaFold-Multimer, on the 17 heterodimer Benchmark 2 dataset from [2]

| Target | DockQ Score |
| --- | --- |
| 5ZNG | 0.69 |
| 6A6I | 0.02 |
| 6GS2 | 0.51 |
| 6H4B | 0.76 |
| 6IF2 | 0.68 |
| 6II6 | 0.83 |
| 6ONO | 0.63 |
| 6PNQ | 0.41 |
| 6Q76 | 0.91 |
| 6U08 | 0.96 |
| 6ZBK | 0.76 |
| 7AYE | 0.85 |
| 7D2T | 0.86 |
| 7M5F | 0.85 |
| 7N10 | 0.84 |
| 7NLJ | 0.06 |
| 7P8K | 0.05 |

**Table S3.** Performance on the *Recent-PDB-Multimers* dataset, evaluated on homology-reduced chain pairs with low training set similarity (see subsection 7.8), broken down into DockQ categories as in Table S1.

| Method | Mean DockQ Score | Incorrect % | Acceptable % | Medium % | High % |
| --- | --- | --- | --- | --- | --- |
| AlphaFold-Linker | 0.476 | 35.7 | 10.1 | 25.5 | 28.7 |
| AlphaFold-Multimer (5x1) | 0.534 | 29.1 | 10.6 | 25.0 | 35.3 |
| AlphaFold-Multimer (5x5) | 0.541 | 27.6 | 11.5 | 24.8 | 36.1 |

| Method | Mean DockQ Score | Incorrect % | Acceptable % | Medium % | High % |
| --- | --- | --- | --- | --- | --- |
| AlphaFold-Linker | 0.290 | 57.6 | 10.4 | 20.6 | 11.4 |
| AlphaFold-Multimer (5x1) | 0.492 | 31.4 | 12.2 | 32.0 | 24.4 |
| AlphaFold-Multimer (5x5) | 0.504 | 30.4 | 11.3 | 32.7 | 25.5 |

## Notes

### Competing Interest Statement

The authors have declared no competing interest.

### Summary of Updates

This revision updates the results for new models trained with a between chain centre-of-mass loss, re-weighting of the violation losses and removal of the prokaryote specific MSA pairing. These changes significantly reduce the number of structures with clashes and improve the overall accuracy.

